# Physical constraints on the social-genetic dynamics of mtDNA inside plant mitochondria

**DOI:** 10.64898/2026.11.19.767932

**Authors:** Joanna M. Chustecki, Andrew J. Summers, Jie Wang, Hugh G. Dickinson, Iain G. Johnston

## Abstract

Mitochondrial DNA (mtDNA) encodes essential cellular machinery and is contained within mitochondria. This physical containment shapes the genetic dynamics of mtDNA populations in the cell. In plants (and other non-bilaterians), mtDNA colocalisation is essential for recombination, while other cellular priorities may favour mtDNA spacing through the cell. Given possible physical interactions between mtDNA and the inner mitochondrial membrane and its cristae, it is not clear how readily or often this colocalisation may occur. Only limited, indirect evidence exists supporting the capacity for plant mtDNAs to physically colocalise. Here, we survey evidence from recent and new single-cell timelapse microscopy to quantitatively characterise the motion of mtDNA nucleoids inside vegetative plant cells. We combine biophysical arguments with new microscopy data and spatial statistics to demonstrate the subdiffusive but not internally static nature of nucleoids, demonstrating that they are enriched at the termini of elongated mitochondria and well-spaced through the cell but capable of colocalisation. We also describe other modes of physical motion that may impact the genetic dynamics of mtDNA populations, and discuss the implication of this physical behaviour for cytoplasmic genetics.

## Introduction

Mitochondrial DNA (mtDNA) is an essential component of mitochondria across most eukaryotes, encoding important mitochondrial genes. Physically, mtDNA is contained in nucleoid structures in the mitochondrial matrix, with some evidence that it is tethered to the inner mitochondrial membrane in humans and yeast (Macuada et al., 2025). The physical location of an mtDNA molecule assigns it a local “sphere of influence”, in which its gene products dominate the machinery in that region (Busch et al., 2014; Jakubke et al., 2021).

In bilaterian animals, mtDNA molecules are typically physically separated, with biophysical mechanisms including “pearling” proposed to support this separation (Landoni et al., 2026). Separation is important for even distribution of nucleoids and faithful partitioning at cell divisions (Glastad & Johnston, 2022; Jajoo et al., 2016). As few processes in bilaterians require two mtDNA molecules to physically meet, there is apparently little downside to this separation.

However, outside bilaterians, there is a ubiquitous process requiring physical interaction between mtDNA molecules. mtDNA recombination occurs commonly, and is essential in several pathways for the repair and segregation of genetic damage, including template repair and segregation via gene conversion (Broz et al., 2022, 2024; Edwards et al., 2021; Preuten et al., 2010; Zwonitzer et al., 2024). These processes are particularly common in plants, with recombination inferred to readily occur on developmental, and cellular, timescales (Broz et al., 2024, 2025).

In plants, mitochondria are typically highly fragmented, with each mitochondrion containing on average less than a complete mtDNA genome (Arimura et al., 2004; Chustecki et al., 2025; Chustecki & Johnston, 2024; Johnston, 2019; Logan, 2010a; Preuten et al., 2010). There is therefore an unexplained physical question accompanying this genetic behaviour. If mtDNAs are isolated in fragmented organelles, and potentially fixed in place even within an organelle, how do they ever meet to under recombination?

(Arimura et al., 2004) and (Chustecki et al., 2025) provide open-access video data of simultaneous mitochondrial and nucleoid visualisation inside plant cells. Previous reported dynamics include the fusion and fission of mitochondria with uneven nucleoid content, consistent with the observation that not all plant mitochondria contain mtDNA (Arimura et al., 2004; Preuten et al., 2010), and that mitochondria that do contain mtDNA move on average faster, cover larger regions of the cell, and encounter other mitochondria more than those without mtDNA (Chustecki et al., 2025). In parallel, theoretical arguments suggest that preferentially supporting fusion of mitochondria containing mtDNA can increase the efficiency of mtDNA repair (Johnston et al., 2026).

Here, we survey the available video data of mitochondria and nucleoids along with new data from *Arabidopsis* and wheat, and couple this data with biophysical and statistical arguments to quantitatively characterise the physical dynamics of mtDNA populations inside plant cells.

## Results

### mtDNA is subdiffusive inside the mitochondrial matrix

We surveyed the video data from (Arimura et al., 2004) and (Chustecki et al., 2025) to characterise the physical behaviour of mtDNA nucleoids throughout the cell. We manually identified case study behaviour and used the established video analysis pipeline (Chustecki et al., 2021) with TrackMate (Tinevez et al., 2017) to analyse mtDNA motion (see Methods). Throughout, we will work with the original raw pixel data, rather than smoothing or transforming, to ensure that our observations do not arise from artefacts of any such processing. Several consistent principles are immediately apparent from these data, including common arrangement of nucleoids in pairs that stay colocalised for extended time periods, and positioning of nucleoids at the termini of elongated mitochondria (Fig. 1A).

**Figure 1.**
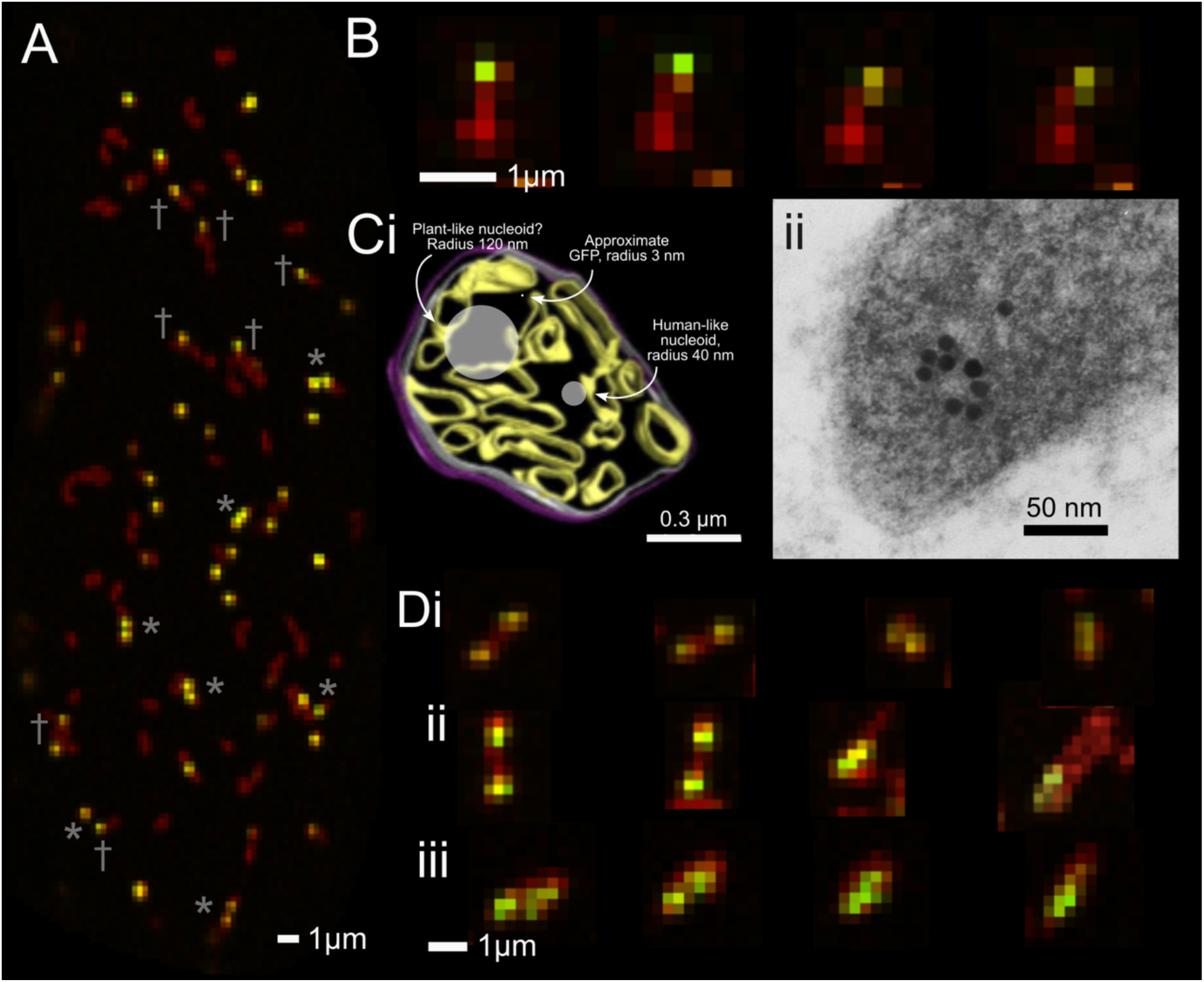
Examples of characteristic nucleoid structure and dynamics in plant epidermal tissue. **(A)** Mitochondria (red) and nucleoids (yellow, from green mtDNA signal and red mitochondrial signal). Many mitochondria are “empty” of mtDNA; nucleoids often appear in persistent pairs (*); in elongated mitochondria, nucleoids often appear at a terminus (†). **(B)** An example of a nucleoid showing no detectable diffusion through the matrix of its mitochondrion, over 300 s. **(C)** (i) Electron microscopy of internal cristae structure of an Arabidopsis mitochondrion from (Golin et al., 2020) (licensed under CC-BY), with approximate sizes of GFP, human mtDNA nucleoids, and estimated plant nucleoids (see Methods) superimposed. (ii) *In situ* nick translation/biotin-gold-labelled mtDNA nucleoid in *Arabidopsis* male germ line (transmission electron microscopy from (Jones et al., 1990), reproduced with permission). **(D)** Examples of observable relative motion of nucleoids within their mitochondria. In each case, contiguous red signal and collective motion of the associated mitochondrial object suggests a single housing organelle, while two initially distinct nucleoid signals detectable move towards overlap.

The first observation we pursue is that the relative positioning of nucleoids inside their mitochondrion is rather stable. An example (Fig. 1B) involves a mitochondrion of length ∼1 μm containing an mtDNA that is at one extreme end. Over the course of 300 seconds there is no noticeable shift in position away from that extreme end, corresponding to no internal motion visible at the resolution limit (Fig. 1B). This specific example is an example of no relative motion over a relatively long period, but many other examples exist where a mitochondrion is imaged over >10 s with no relative mtDNA motion. The microscopy scale is 1 pixel ≈ 0.25 μm. For the absence of an observable movement over a 4 pixel scale (conservatively corresponding to the diameter of the mtDNA punctum), any shift has to be < 1 μm over 300 seconds. If this motion were due to simple diffusion, we would conclude that the diffusion constant for that mtDNA, *D* ≲ △*x*_det_^2^/2*t*, so *D* ≲ 1^2^⁄(2 ⋅ 300) ≈ 0.0017 *μ*m^2^/s.

In the Supplementary Information, we show that even under a collection of conservative biophysical assumptions, we would expect a plant mtDNA nucleoid experiencing free diffusion in the matrix to show at least D ≈ 0.33 μm^2^ s^-1^, orders of magnitude faster than observed. We can therefore conclude that free diffusion in the matrix is not an accurate picture for mtDNA nucleoids. This biophysical reasoning suggests a length scale for a plant mtDNA nucleoid at least on the scale of the crista structure inside the organelle (Fig. 1C), supported by EM images (Fig. 1Cii). It seems therefore likely that, even in the absence of explicit tethering, the internal structure of the IMM provides substantial hurdles to nucleoid motion in the matrix.

### Detectable motion of mtDNA inside mitochondria

Clearly, nucleoids are relatively constrained inside a mitochondrion. However, if any relative motion of nucleoids can occur in the same organelle, the possibility arises for (albeit slow) physical colocalisation and therefore genetic interaction. We next looked for evidence that such relative motion could be possible.

The challenge in characterising internal motion of mtDNA is separating the mtDNA motion from that of its host organelle. If two mtDNAs are in different mitochondria, the relative motion of the two organelles will confound any attempt to measure internal motion of nucleoids (Supp. Fig. 1). We therefore looked for examples fulfilling several criteria: continuous matrix signal linking the two nucleoid puncta through the entire time course of an observation, and correlated motion of both puncta and matrix signal through the course of an observation. Several examples are shown in Fig. 1D. In these cases, a relative distance of at least 0.5 μm is covered over the course of around 30 s, suggesting an effective diffusion rate of *D* ≈ Δ*x*_det_^2^ ⁄2*t* ≈ (0.5)^2^⁄(2 ⋅ 30) ≈ 0.004 *μ*m^2^/s.

If crista structure provides the most substantial physical barrier to nucleoid motion, these may reflect examples of nucleoids “hopping” from one crista-delimited compartment to another. Using these data it is impossible to distinguish motion due to internal diffusion from motion due to conformational change of the organelle (squashing, stretching, twisting; Supp. Fig. 1). However, whatever the underlying dynamic, these observations suggest that mtDNAs can indeed move relative to each other, inside an organelle.

However, these events are relatively rare. While the resolution of the images makes a precise estimate impossible, we estimate that around 1% of elongated mitochondria containing at least two mtDNA puncta showed such detectable internal motion, over the several minutes of imaging of each case. Roughly, 1-10% of nucleoids on average are present in a mitochondrion containing at least two nucleoids (hence, with a potentially accessible “partner” for interaction).

### Physical properties of nucleoid and mitochondrial populations

We next asked about fundamental physical properties of the mitochondrial and nucleoid populations in the cell: number, density, speed, and separation (Fig. 2A-C). These quantities have been reported before (Chustecki et al., 2021, 2025; Logan, 2003, 2010b); we include them here for completeness and context in this study, and to support BioNumbers-style quantitative cell biological calculations (Milo et al., 2010; Phillips & Milo, 2009).

**Figure 2.**
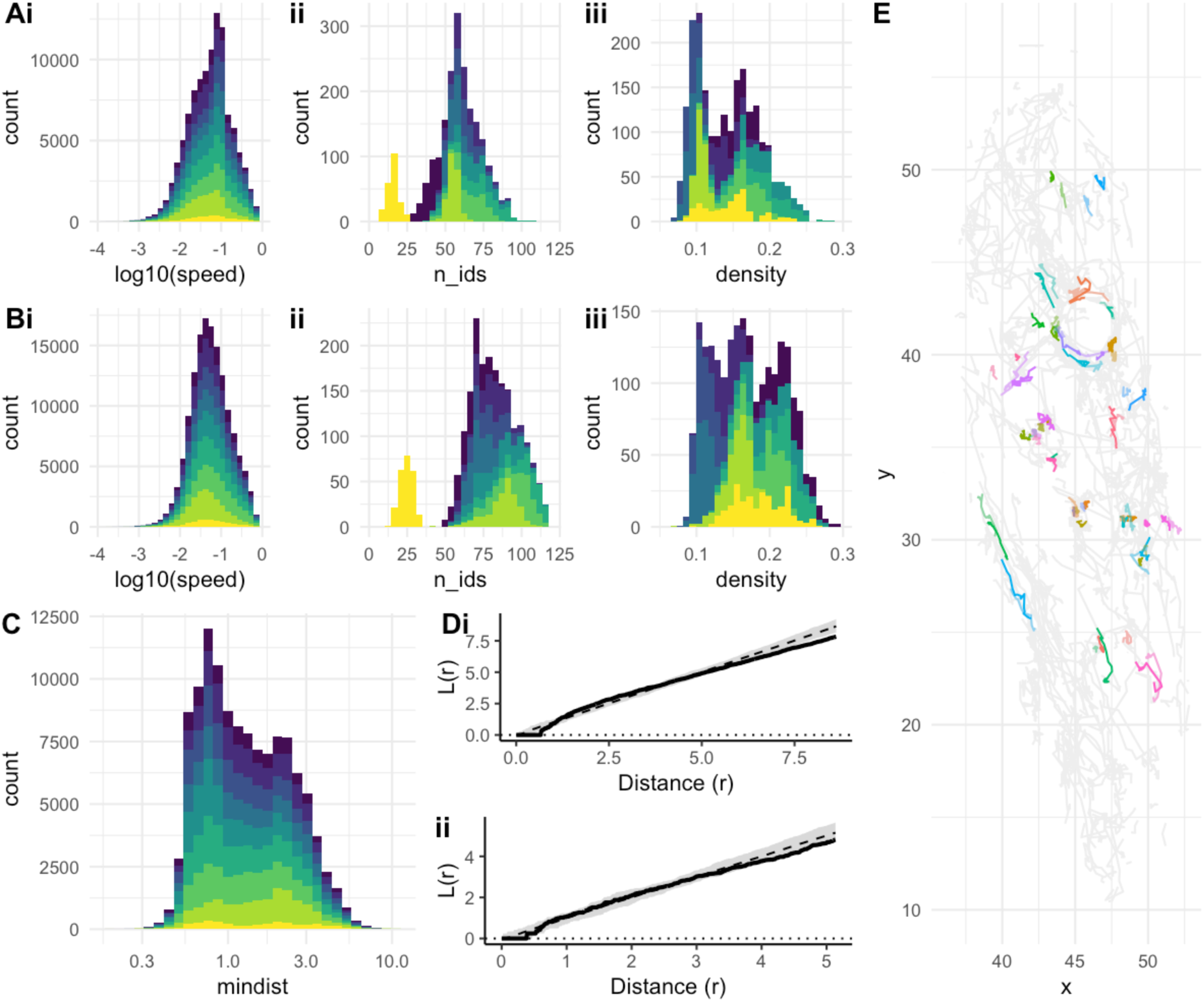
Physical behaviour of of mitochondria and nucleoids. **(A-B)** (i) Speed, (ii) number of puncta, and (iii) density of puncta for (A) nucleoids, (B) mitochondria. Different colours give statistics from different independent biological samples. **(C)** Nucleoid distance to nearest neighbour. **(D)** Ripley’s L-statistic L(r), reporting the average number of neighbours within distance r scaled by overall density. (i) Mitochondria; (ii) nucleoids. Black line shows example L(r) from real cell data; grey intervals show behaviour of L(r) under a null hypothesis of random distribution. **(E)** Persistent pairs. The set of nucleoids trajectories observed in a cell (grey). Persistent pairs, defined as two nucleoids remaining within 2μm of each other for at least 40s, are coloured, with one member darker and one lighter.

Nucleoids are typically present at a density around 0.1-0.2 μm^-2^, with between 25-100 typically visible in one frame of the source data. Assuming a roughly cuboidal structure for the cell, this suggests between 100-500 nucleoids per cell (accounting for the different faces of the cuboid, with the vacuole occupying the centre) – compatible with the whole-cell observations in (Chustecki et al., 2025). Instantaneous speeds are commonly between 0.01-0.1 μm s^-1^ and never exceed 1 μm s^-1^. Minimum distances to the closest nucleoid follow a bimodal distribution, with peaks often around 1 μm and 2.5 μm, with distances greater than 5 μm very rare. Overall mitochondrial speed distributions occupy a similar range with a slight shift to lower values, consistent with the observation that organelles with mtDNA move slightly faster than those without (Chustecki et al., 2025). There are typically roughly two times as many mitochondrial puncta as nucleoid puncta, consistent with the observation of “empty” mitochondria, but suggesting a higher proportion of mtDNA-containing mitochondria than the 1/3 previously observed; however, this discrepancy is unsurprising given previously observed cell-to-cell heterogeneity in the “empty” proportion of mitochondria (Chustecki et al., 2025). The minimum distance between a nucleoid and its nearest neighbour (which is often in a different mitochondrion) is somewhat bimodally distributed across a range of cell sizes and geometries (Fig. 2C), with modal distances both <1 μm and around 2 μm, corresponding to common near adjacency between nucleoids (Fig. 1A).

### mtDNA nucleoids form persistent pairs in the cellular population

Following this near adjacency, we next pursued the observation that mtDNA nucleoids often colocalise in “persistent pairs”, where two nucleoids remain within a distance <2 μm of each other for a period of many seconds to minutes, often traversing a considerable distance in the cell. Fig. 1A gives a snapshot example of persistent pairs; Fig. 2E illustrates some examples over time. Over 9 independent sample videos analysed, 10.4 ± 2.7% of nucleoids were part of a persistent pair at any given time.

In many cases, this colocalisation is associated with two separate mitochondria, which colocalise and may transiently fuse. It is known from (Arimura et al., 2004) that fusion mixes matrix content on a timescale of a few seconds, but the physical separation of nucleoids persists in the majority of these colocalisation cases. Hence, even if not all long-term colocalisations result in fusion, we can infer that mtDNA nucleoids are not mixed as readily as other matrix contents on fusion events. Slow nucleoid spread is compatible with observations in protoplast fusion (Sheahan et al., 2005) where nucleoids were observed to spread through the cellular population of organelles on a timescale around 72 hours, compared to the 24 hour mixing time of other mitochondrial content (itself rather longer than the ∼2 hour timescale observed in onion epidermis (Arimura et al., 2004)).

### Spacing of mitochondria and nucleoids through the cell

It has been hypothesised that mitochondria face a tension between physical proximity (for exchange of contents) and physical spacing (to ensure an even distribution throughout the cell) (Chustecki et al., 2021; Chustecki & Johnston, 2024, 2026). To explore the spacing of mitochondria and mtDNA throughout the cell, we used spatial statistics – specifically, Ripley’s L-function, which measures the number of neighbours a point is expected to have within a given distance compared to a null hypothesis of random spread. Fig. 2D shows example comparisons of observed neighbour density against this null hypothesis, demonstrating that both mitochondria and nucleoids demonstrate behaviour very compatible with maintaining uniform spread throughout the cell (full results in Supp. Fig. 2). Despite the considerable clustering and pairing behaviour observed, the overall populations of nucleoids and organelles are statistically indistinguishable from a random spread through the cell.

### mtDNA is enriched at termini of elongated mitochondria

Next, mtDNA nucleoids seem to be enriched at the extreme ends of elongated mitochondria (Fig. 1A), also observed in (Arimura et al., 2004). To test this hypothesis quantitatively, we manually identified instances of two cases: (i) where a nucleoid signal appears at the end of an elongated mitochondrion, with no discernible matrix signal beyond it; and (ii) where a nucleoid signal appears inside an elongated mitochondrion, with matrix signal on both sides. The probability that a circular signal of radius r (the nucleoid punctum), placed with its center randomly on a line of length L (the mitochondrion) obscures one end of the line is *p*_0_ = *min*(1, 2*r*/*L*), giving the null probability of observing case (i) if nucleoids are randomly placed inside mitochondria. Over 42 cases, with r = 1.5 px and L ≥ 6 px, we manually observed 29 instances of (i) and 13 instances of (ii) (some examples in Fig. 3A, others in Supp. Fig. 3). A binomial test against the null hypothesis of uniform distribution, with a conservative L = 6 px (neglecting the higher power associated with longer mitochondria) and thus a *p*_0_ = 0.5, gave p = 0.020, illustrating an enrichment of nucleoid positioning towards the ends of elongated mitochondria.

**Figure 3.**
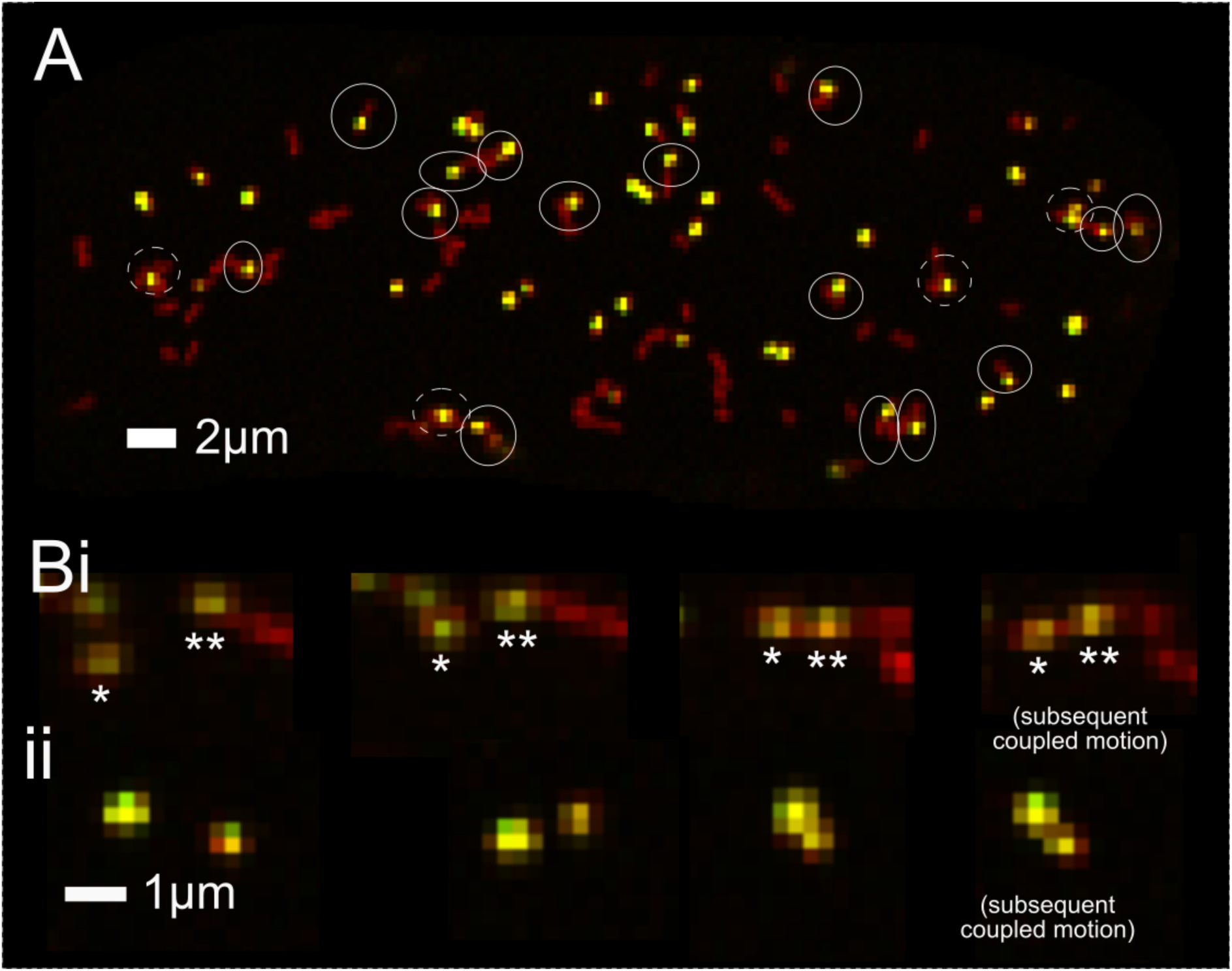
Nucleoid position within and between mitochondria. **(A)** Example instances of nucleoid positions in elongated mitochondria. Solid circles ring terminal nucleoids; dashed circles ring nucleoids not at the terminus of their mitochondrion. **(B)** (i)-(ii) Candidate fusion events supporting the colocalisation of mtDNA nucleoids. The asterisks in (i) track the identity of specific nucleoid signals of interest.

### Exchange of mtDNA between organelles via fission-fusion

Previous work exploiting mixed fluorophores has characterised fission-fusion dynamics in plant mitochondria in detail (Arimura, 2018; Arimura et al., 2004). Without mixed fluorophores, our ability to confirm genuine fusion events is limited, but we do observe several candidate events where two initially separate mitochondria colocalise to form a continuous matrix signal which continues to move consistently throughout the data series. In several cases, this potential fusion leads to a colocalisation of previous separate mtDNA puncta (Fig. 3B).

One question is how readily mtDNA nucleoids can be transferred from one mitochondrion to another during a fusion-fission cycle. This impacts the potential for template repair (Johnston et al., 2026) and mtDNA distribution through the cell (Arimura et al., 2004; Sheahan et al., 2005). The physical constraint of nucleoids within their organelles has a strong impact on how nucleoids are “inherited” by the daughter mitochondria following a fission event. We consider the following model picture to analyse this impact. A nucleoid is present in a spherical mitochondrion, which fuses at a given point with another spherical mitochondrion that lacks a nucleoid. The product of this fusion is an elongated mitochondrion containing the nucleoid. This elongated mitochondrion then undergoes fission. Which daughter mitochondrion now contains the nucleoid?

The answer depends on the original position of the nucleoid and the geometry of the fission plane. In the Supplementary Information we analyse a simple 2D representation of this model. We picture the nucleoid present at an angular position *θ* relative to the point of fusion, with the fused mitochondrion being elliptical with semi-major and semi-minor axis lengths *a* and *b*. The crucial quantity is then the new position of the nucleoid within the fused mitochondrion, which can be represented as a parametric angle *θ*′, the solution to an elliptic integral equation (given, including a possible simplification, in the Supplementary Information). We consider two circumstances for fission. In the first, the fission plane goes through the centre of the elongated mitochondrion, at an angle *φ* to the fusion plane that formed it. In this case, the nucleoid shifts if *φ* > tan^−1^(*a* tan *θ*′ / *b*). In the second case, the fission plane runs parallel to the fusion plane at a distance *d* from its centre, and the nucleoid shifts if *d* > *a* sin *θ*′. Both these conditions make it challenging, but not impossible, for a tightly constrained nucleoid to switch host organelle through a fission-fusion cycle (Fig. 4), particularly if nucleoids are enriched at the termini of elongated mitochondria.

**Figure 4.**
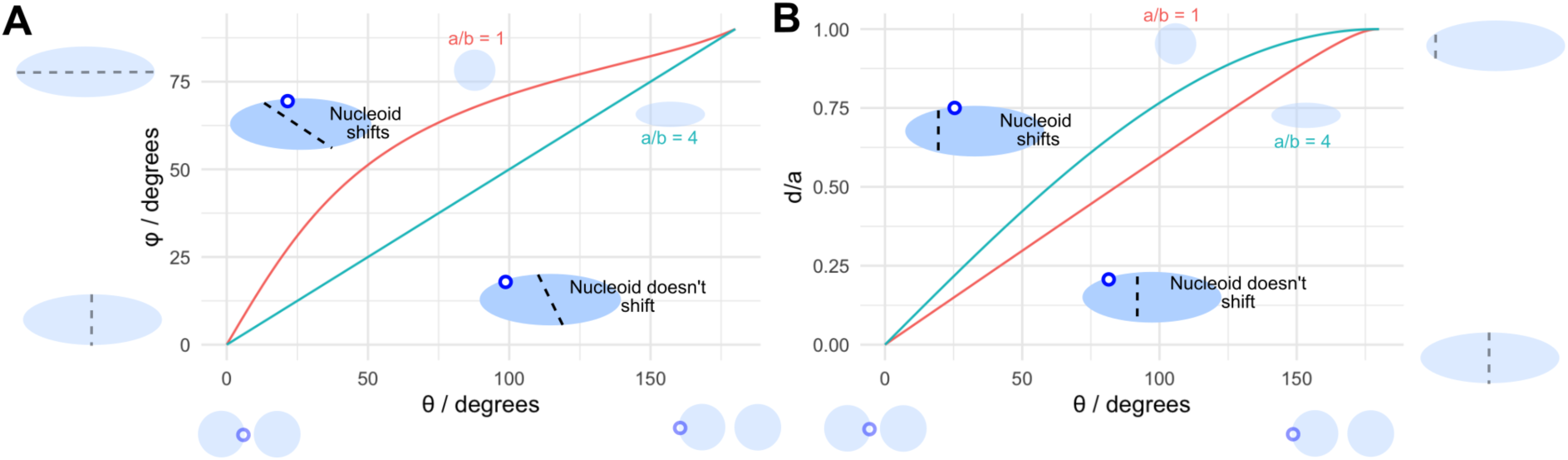
Conditions for exchange of a nucleoid through a fusion-fission cycle. Two model mitochondria fuse at a point to form an elongated mitochondrion with preserved membrane content. The left-hand mitochondrion contains a nucleoid. Fission subsequently fragments this elongated mitochondrion. On the horizontal axis, *θ* gives the initial angular position of the nucleoid relative to the fusion point, before fusion takes place. **(A)** The fission plane goes through the centre of the elongated mitochondrion, at angle *φ* to the fusion plane (vertical axis). **(B)** The fission plane runs parallel to the fusion plane, at a proportional distance d/a between the centre of the elongated mitochondrion and one terminus. The lines in each case show the conditions on *θ* and fission plane geometry for the nucleoid to “shift” from the left-hand original mitochondrion to the right-hand daughter. The two lines give results for fused mitochondria of different aspect ratios a/b.

Documented fission events in plant mitochondria with nucleoids involve fission case (ii) with the nucleoid around a terminus of the elongated mitochondrion (implying *θ* around 180°), and d/a roughly 0.3-0.5 (Fig. 4C in (Arimura et al., 2004)) or *φ* < 45° (Fig. 4B in this manuscript). Both these cases fall firmly in the regions of Fig. 4 where the nucleoid does not shift host mitochondrion.

### Consistency across species and tissues

All the quantitative observations above have come from hypocotyl epidermis in *Arabidopsis thaliana* seedlings. To assess how general these principles might be across plants and tissues, we established a joint mitochondrial/mtDNA imaging protocol in wheat seedling roots (see Methods).

In this tissue, cells are smaller and often less elongated, and substantial transvacuolar strands often convey cytoplasm and mitochondria through the vacuole at the centre of the cell. However, mitochondrial populations in planes of cytoplasm share notable similarities with the situation in Arabidopsis hypocotyl (Fig. 4). Roughly half of mitochondria have no associated mtDNA signal; nucleoid signals are enriched towards the termini of elongated mitochondria and there is some evidence of persistent pair behaviour (Fig. 5A). Also, while nucleoids again appear relatively static within their host organelle (suggesting a similar bound on diffusive motion as observed in Fig. 1B), some relative motion of nucleoid signals inside an organelle is observed, along with relative motion of nucleoids due to deformation of the host organelle, and fission followed by subsequent decoupling (Fig. 5B).

**Figure 5.**
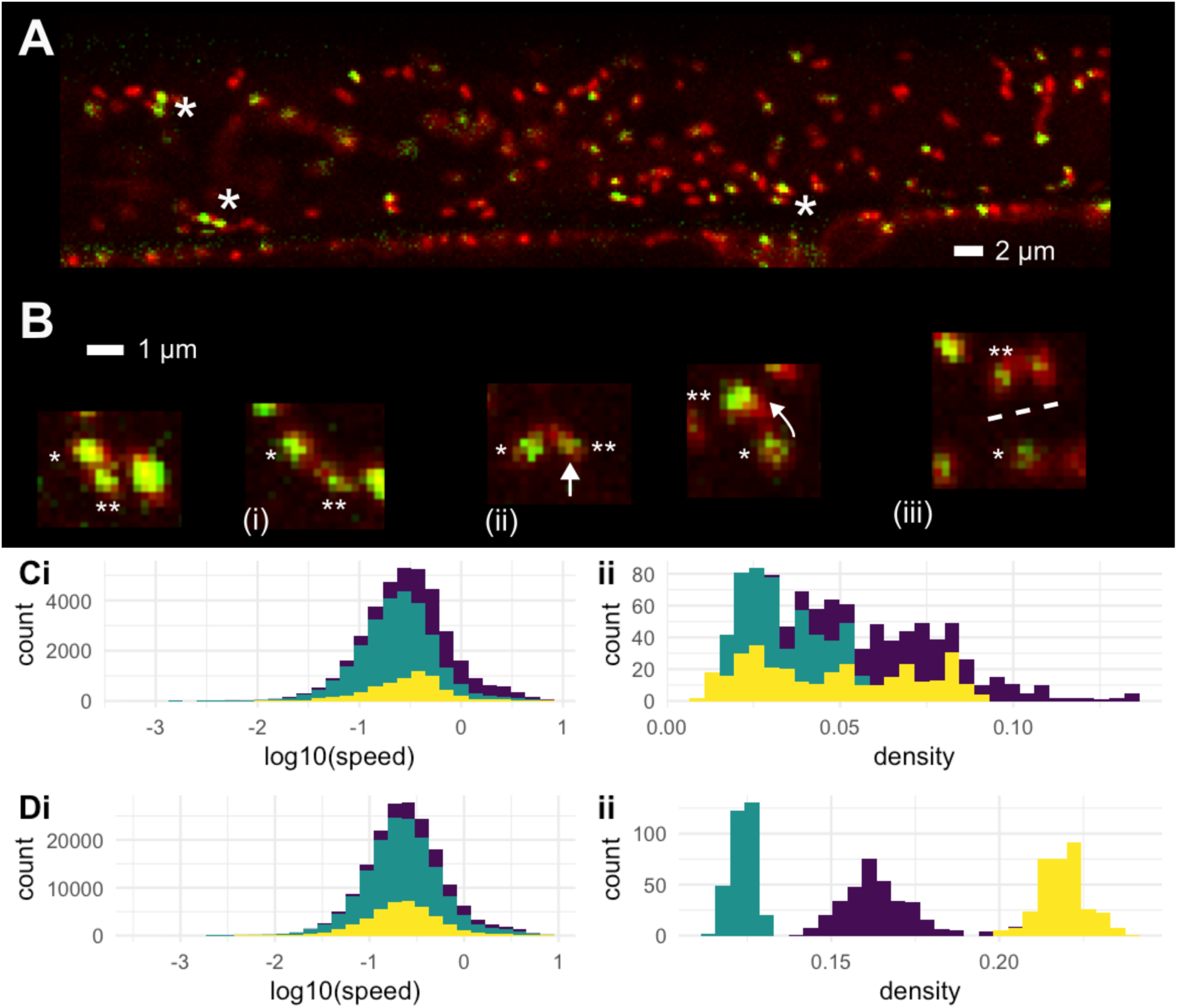
Mitochondria and mtDNA nucleoids in wheat root epidermis. **(A)** Overview of an epidermal cell. Mitochondria are stained with MitoTracker Deep Red (red) and nucleoids stained with SyBR Green (green, for a yellow overlap). Persistent pairs (*) and a limited number of mitochondria containing mtDNA are observed. **(B)** A single event from (A) demonstrating several of the dynamic features observed in Arabidopsis. One nucleoid (*) stays constrained at the end of its organelle over time. The other (**) moves relative to the first without organelle deformation, moves relatively as the organelle subsequently deforms through bending, then decouples via fission and moves independently. **(C)** Speed (i) and density (ii) distributions of nucleoid signals across wheat cells. Different colours give different independent biological samples. **(D)** Speed (i) and density (ii) distributions of mitochondrial signals across wheat cells.

The physical properties of the mitochondrial system here appear similar to the *Arabidopsis* case (Fig. 5C-D). Instantaneous speeds of both mitochondria and nucleoids are typically between 0.05-0.5 μm s^-1^, distributed similarly to those in *Arabidopsis* hypocotyl but shifted faster. Observed nucleoid densities are lower than *Arabidopsis*, around 0.02-0.1 μm^-2^, with roughly 1/3 of mitochondria containing detectable nucleoids. Overall mitochondrial density was more similar to *Arabidopsis*, but differed substantially between samples (Fig. 5Dii).

## Discussion

Taken together, this data paints a picture of nucleoids that are physically constrained within their host mitochondria, but not completely prevented from moving internally. Nucleoids throughout the cell are well-spaced as a population, but individually are often closely colocalised with a nearby partner, likely following, during, or immediately prior to fusion of the host organelles. This picture is compatible with a model where mtDNA colocalisation is relatively rare, at least in the vegetative tissue types we observe here. As a very rough estimate from *Arabidopsis*, as roughly 1% of nucleoids in elongated mitochondria demonstrated colocalisation capacity through ∼ 180 s of imaging, and 1-10% of nucleoids are typically present in elongated mitochondria, an estimated rate for colocalisation capacity would be 0.002-0.02 events per nucleoid per hour. For comparison, (Broz et al., 2024) gave a not inconsistent, but also very indirect, estimate of 0.1 gene conversion events per mtDNA per cell division during plant development, based on genetic observables. However, it is impossible for now to more directly tie the physical and genetic observations, which in any case are in different tissues and cellular populations.

Is plant mtDNA tethered to the inner membrane? Characterisation of DNA- and membrane-associated factors in *Arabidopsis* mitochondria have revealed some candidates that could be involved in explicit tethering (Kim et al., 2021a), but conclusive molecular answer to our knowledge remains absent. The physical data in this study cannot conclusively answer this, although none of the observations we have made require tethering, and our observation of within-mitochondrion nucleoid motion is hard to reconcile with a picture of complete tethering. Rather, the steric hindrance provided to bulky nucleoids by cristae – likely also a feature in other species (Dua et al., 2026) -- would seem sufficient to explain the general constraint and occasional motion we observe within mitochondria.

Our picture of physical motion within the organelle, restricted by inner mitochondrial membrane (IMM) structure but not permanent tethering, suggests several testable hypotheses. If IMM structure is mainly responsible for nucleoid constraint, mutating cristae structure (for example, through a MICOS mutant (Li et al., 2019)) should allow freer mtDNA diffusion -- whereas if tethering is the main constraint, such mutants should have limited impact on mtDNA motion. A further hypothesis builds upon the idea that the rare, highly-fused mitochondrial state seen in the shoot apical meristem (SAM) may support mtDNA recombination (Edwards et al., 2021; Segui-Simarro et al., 2008). If large-scale fusion indeed serves to support mtDNA colocalisation and recombination, segregation of mtDNA heteroplasmy should be challenged in fusion mutants – a prediction that could be tested in artificially heteroplasmic lines (Broz et al., 2022, 2024). By contrast, if the role of large-scale fusion is mainly bioenergetic (Hoitzing et al., 2015) or to support the even physical partitioning of mtDNA between daughters at cell divisions (Glastad & Johnston, 2022; Jajoo et al., 2016), segregation should not be impacted by compromised fusion limiting recombination capacity.

Across plant species, and tissues within a species, mitochondrial dynamics vary dramatically (Logan, 2010b; Logan & Leaver, 2000; Oldenburg & Bendich, 2015). In addition to the SAM behaviour above, large-scale *Arabidopsis* mitochondrial fusion is also observed post-imbibation (Paszkiewicz et al., 2017). Filamentous mitochondria and dynamics associated with uneven inheritance at divisions are observed in zygotes and related cell types in *Arabidopsis* and *Pelargonium* (Kimata et al., 2020; Kuroiwa et al., 2002). *Arabidopsis* mitochondria in cells that are packed with chloroplasts move in the highly confined regions available between these larger organelles (Oikawa et al., 2021); mitochondria in pollen tubes move rapidly in highly constrained physical domains (Liu et al., 2024). Cross-species diversity is also notable, with *Physcomitrella* mitochondria moving around 1/10 the speed of those observed in *Arabidopsis* (Mueller & Reski, 2015). Our goal here is to explore the motion of nucleoids on the background of overall organelle motion – here, the comparison between *Arabidopsis* hypocotyl and wheat root epidermis showed general qualitative consistencies alongside quantitative differences in physical motion. A comprehensive picture of coupled mitochondrial and mtDNA dynamics across species and tissues must remain a future target.

As many mitochondria in plant cells lack mtDNA, recent modelling studies have considered how individual mitochondria access the machinery they require (Giannakis et al., 2022) and how mtDNA maintenance through recombination-mediated processes may take place (Johnston et al., 2026). These models have suggested that the exchange of more freely-diffusing gene products (mRNA, and/or mobile proteins) through fusion-fission cycles can accomplish complementation and maintenance – but that exchange of mtDNA through fusion-fission increases the efficacy of these processes. Our observations here support the general picture that mtDNA is tightly constrained within mitochondria. However, our observations are very compatible with rare and/or slow events that can support mtDNA mixing through the population – slow diffusion within multi-nucleoid mitochondria, and “shifting” of nucleoids between fission products through geometry. Longer-term joint observations of live-cell mitochondria and nucleoid dynamics will provide better characterisation of these rare events, and thereby help characterise the existence and impact of mtDNA exchange through the “social network” of plant mitochondria in the cell.

## Methods

### Biophysical modelling of diffusion, fission, and fusion

In the Supplementary Information we describe the physical and geometric arguments supporting our discussion of mtDNA nucleoid constraint in the mitochondrial matrix, and the possibility of mitochondria exchanging constrained nucleoids through fission-fusion cycles.

### Video analysis

Given video data with mitochondrial matrix and nucleoid signal in different channels, we split the channels and analysed both with TrackMate (Tinevez et al., 2017). We used a Laplacian of Gaussians (LoG) point detector with subpixel localisation but not median filter preprocessing (which tended to blur together distinct but adjacent puncta), using characteristic diameters of 0.7 μm for mitochondria and 0.4 μm for nucleoids. Ǫuality thresholds were chosen manually for each sample, so that detected points visually matched distinct signals. We used the simple Linear Assignment Problem (LAP) tracker, with linking maximum distance 2 μm, gap-closing maximum distance 3 μm, and gap-closing maximum frame gap of 2. CSV files describing tracked trajectories were exported and analysed in the R codebase.

### Mitochondrial and mtDNA imaging in Arabidopsis

For imaging of live hypocotyl cells, 5-days old seedlings grown on plates were used. Tissues were mounted in water on glass slides, with double-sided sticky tape as mechanical stress buffer and a glass cover slip.

All laser scanning confocal micrographs with the exception of the 14 mito-mCherry AOX-mtHINESS time lapse videos were taken on an upright Nikon A1-NiE system (RRID:SCR_020318) using NIS-Elements Confocal image acquisition and analysis software, using a 60x water objective. Images were taken using a sequential channel series setting, to minimise cross-channel signal, and the channels used were: GFP, Ex: 488nm, Em: 525 (500-550), TxRed, Ex 561, Em: 595 (570-620), Cy5, Ex: 640nm, Em: 700 (663-738). All time-series images taken on the Nikon system have a frame time of 2.21s.

The 14 mito-mCherry AOX-mtHINESS time-lapse videos analysed were of single hypocotyl cells of 5-day seedlings, taken on an inverted Zeiss LSM 980 with ZEN blue image acquisition and analysis software, using a 63x oil immersion lens. Time lapses were taken in two tracks, GFP, Ex: 488nm, Em: 509 nm (490-561nm), mCherry, also capturing chloroplast autofluorescence, Ex: 587, Em: 610nm (596-685nm). This collection of time series has a frame time of 2.52 secs.

### Mitochondrial and mtDNA imaging in wheat roots

Wheat seedlings were obtained commercially (Mat og Hage, Norway). Seedlings were surface sterilised with 30% domestic bleach for 1 minute and rinsed three times then germinated on wet cotton wool under natural day/night cycles. At 6 days the seedlings were incubated for 30 minutes with 0.2 μM MitoTracker Deep Red (ThermoFisher Scientific, M22426) and a 1:100000 dilution of SYBR Green I (ThermoFisher Scientific, S7563) based on the protocol from (Kim et al., 2021b), then placed on an imaging dish in a drop of water. Imaging was done using an Olympus FV3000 point-scanning confocal microscope. For SYBR Green I, excitation was detected by a laser at 488 nm and emission collected at 500–540 nm, and for MitoTracker Deep Red FM, excitation was detected with a 640 nm laser and emission collected at 650–750 nm, all at 60x magnification.

### Statistical analysis

Statistical analysis and visualisation was done in R (R Core Team, 2022). We used dplyr (Wickham et al., 2023) for data manipulation, ggplot2 (Wickham, 2016), ggpubr (Kassambara, 2020), and viridis (Garnier et al., 2021) for data visualisation, and spatstat.geom and spatstat.explore (Baddeley et al., 2016) for spatial statistics.

## Data and code availability

All new video data, code, CSV datafiles from the tracking steps, and a directory of particular data used in the case studies in this manuscript is freely available at https://github.com/StochasticBiology/mtdna-motion.

## Acknowledgements

This project has received funding from the European Research Council (ERC) under the European Union’s Horizon 2020 Research and Innovation Programme [Grant agreement No. 805046 (EvoConBiO) to I.G.J.]. This work was supported by the Research Council of Norway through the FRIPRO program [project numbers 357812 and 360449 (MitoPhyto) to I.G.J.]. J.M.C. was financially supported by a WiRe fellowship from the University of Münster Internationalization Fund. I.G.J. thanks the Rudolf Peierls Centre for Theoretical Physics at the University of Oxford for hosting a sabbatical visit during work on this project, and L. Meltzers Universitetsstiftelse from the University of Bergen for supporting this visit.

## Supplementary Information

### Biophysical bounds on mtDNA diffusion inside the matrix

A baseline estimate for diffusion rate in the mitochondrial matrix is given by FRAP measurements of matrix-targeted GFP (Partikian et al., 1998), giving a range of effective diffusion coefficients ∼20–30 μm²/s, with high mobile fractions (>90%). At least for small, soluble proteins, this suggests a picture of the matrix as a continuous compartment which can be explored by diffusion, albeit at around a 3x lower rate than in water [Bionumber 100301 (Milo et al., 2010); (Swaminathan et al., 1997)]. This decrease in motility is likely due to molecular crowding and the cristal structure of the inner mitochondrial membrane.

How can we relate this GFP picture to mtDNA nucleoids? This is impossible without some assumptions about the physical structure of a nucleoid. Electron microscopy in animal cells suggests nucleoids have a radius around 40-50 nm (Kukat et al., 2011; Prachař, 2016). We are not aware of direct imaging of intact nucleoids at the nanometre scale in plant mitochondria, and must be cautious about straightforwardly using animal values, as the plant mtDNA genome is longer, split into heterogeneous molecules, and packaged differently. In *Arabidopsis*, WHIRLY2-dependent nucleoid organization and TEM studies of wild-type and mutant mitochondria show discrete nucleoid-associated electron-lucent regions and punctate structures consistent with compact nucleoprotein assemblies rather than extended DNA coils (Golin et al., 2020). Assuming at least some packaging, we can then bound the size of an *Arabidopsis* nucleoid with the following argument. Human mtDNA is 16.6 kb at 0.34 nm per base pair, for a length of L = 5.6 μm. DNA persistence length is around l_p_ = 50 nm [Bionumber 103112 (Milo et al., 2010); (Garcia et al., 2007)]. The radius of gyration of a worm-like coil is 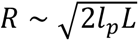 = 750 nm. Packaging therefore decreases the nucleus radius approximately 20-fold to the free polymer estimate. *L* = 125*μm* for the full Arabidopsis mtDNA in one molecule, for R ∼ 3500 nm, which with a 20-fold packaging reduction would give a radius of 180 nm. We can therefore picture a reasonable range for plant mtDNA nucleoid size going from 40 nm (human case, short mtDNA) to 180 nm (estimated complete genome).

If we represent a nucleoid as a compact structure, much larger but otherwise comparable to a protein, we can use the GFP result to estimate expected matrix diffusion rates of a nucleoid. We can picture GFP as a ∼3 nm hydrodynamic tracer. The Stokes–Einstein relation gives D ∝ 1/R. So for the two limits of mtDNA nucleoid size we would expect

D_small_ ∼ 20 / (40/3) = 1.5 μm^2^ s^-1^

D_big_ ∼ 20 / (180/3) = 0.33 μm^2^ s^-1^.

This estimate represents purely size-dependent hydrodynamic slowing within the same crowded environment defined experimentally by GFP FRAP. However, there is a major difference between the length scales of GFP and estimated nucleoids: the internal structure of the mitochondrion involved cristae on the scale of ∼50 nm. While GFP diffusion may be somewhat insensitive to this structure, the motion of larger objects will certaintly be impacted by cristae (see Fig. 1).

### Geometry of fission-fusion events

We model the two mitochondria prior to fusion as circles of equal radius *R*, and they will become an ellipse 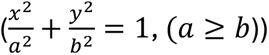. Under the conservation of membrane perimeter, the total perimeter satisfies:

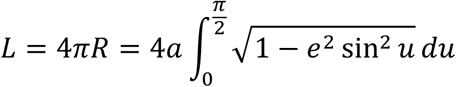

where 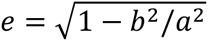 denotes the eccentricity.

Prior to fusion, we write *θ* for the angle at the centre of the nucleoid-containing mitochondria between the location of the nucleoid and the fusion point (Supp. Fig. 4A). The arc length around the circumference to the nucleoid from the fusion point is *s* = *θR*, with *θ* ∈ [0, π] constrained by symmetry.

We model fusion as involving a deformation of each circle to a half-ellipse, with the fusion point expanding to a line interface between the two half-ellipses. This models a 2D projection of a fusion point in 3D expanding to form a circular annulus linking two half-ellipsoidal segments. We assume that membrane content is preserved through the fusion process, so that mathematically, the perimeter of a half-ellipse is equal to the circumference of the original circle. Then we map the original nucleoid position onto the elliptical boundary by preserving its arc length, yielding the parametric relationship:

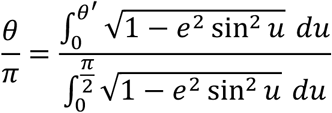

where *θ*′ is the updated parametric angle of the nucleoid on the half-ellipse boundary. This expression, for *θ*′ in terms of *θ*, generally requires a numerical solution of the corresponding integral equation; if *e*^2^ sin^2^ *u* is small, we can apply a first-order Taylor expansion 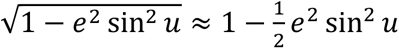, to simplify:

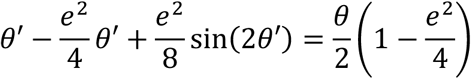

Based on the parametric angle *θ*′, we can derive the geometric angle *ψ* (Supp. Fig. 4B), which is the angle between the fusion plane and the line joining the centre to the nucleoid, with tan 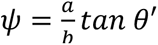.

Without loss of symmetry, we can picture the nucleoid as starting in the “left-hand” mitochondrion. Then the Cartesian coordinates of the nucleoid on the ellipse, if the origin is at the centre, are:

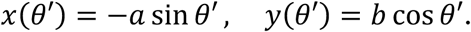

Following fusion, we consider fission as a transverse division that separates the elongated mitochondrion into two smaller mitochondria. We work with two pictures. First (case (i)), we assume that the plane of fission runs through the centre of the elliptical mitochondrion, orientated at angle *φ* relative to the fusion plane (Supp. Fig. 4Ci). In order for the nucleoid to “switch” mitochondria, we require it to lie on the opposite side of the fission plane than it did of the fusion plane. Without loss of symmetry, we can picture the nucleoid as initially in the “left-hand” mitochondrion, and state this requirement as

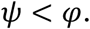

Next (case (ii)), we consider the case where the fission plane is aligned with the fusion plane, but offset by a distance *d* along the major axis of the elongated mitochondrion (Supp. Fig. 4Cii). Again, the nucleoid will “shift” mitochondria if it lies on the opposite side of this fission plane than it did of the fusion plane. For a nucleoid initially in the “left-hand” mitochondrion, the condition for this shift is simply *x*(*θ*′) > −*d*, or

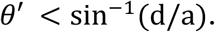

Documented fission events in plant mitochondria with nucleoids involve fission case (ii) with d/a roughly 0.3-0.5 (Fig. 4C in (Arimura et al., 2004)) and an example more like fission case (i) with *φ* < *π*/4 (Fig. 4B in this manuscript).

---

**Supplementary Figure 1.**
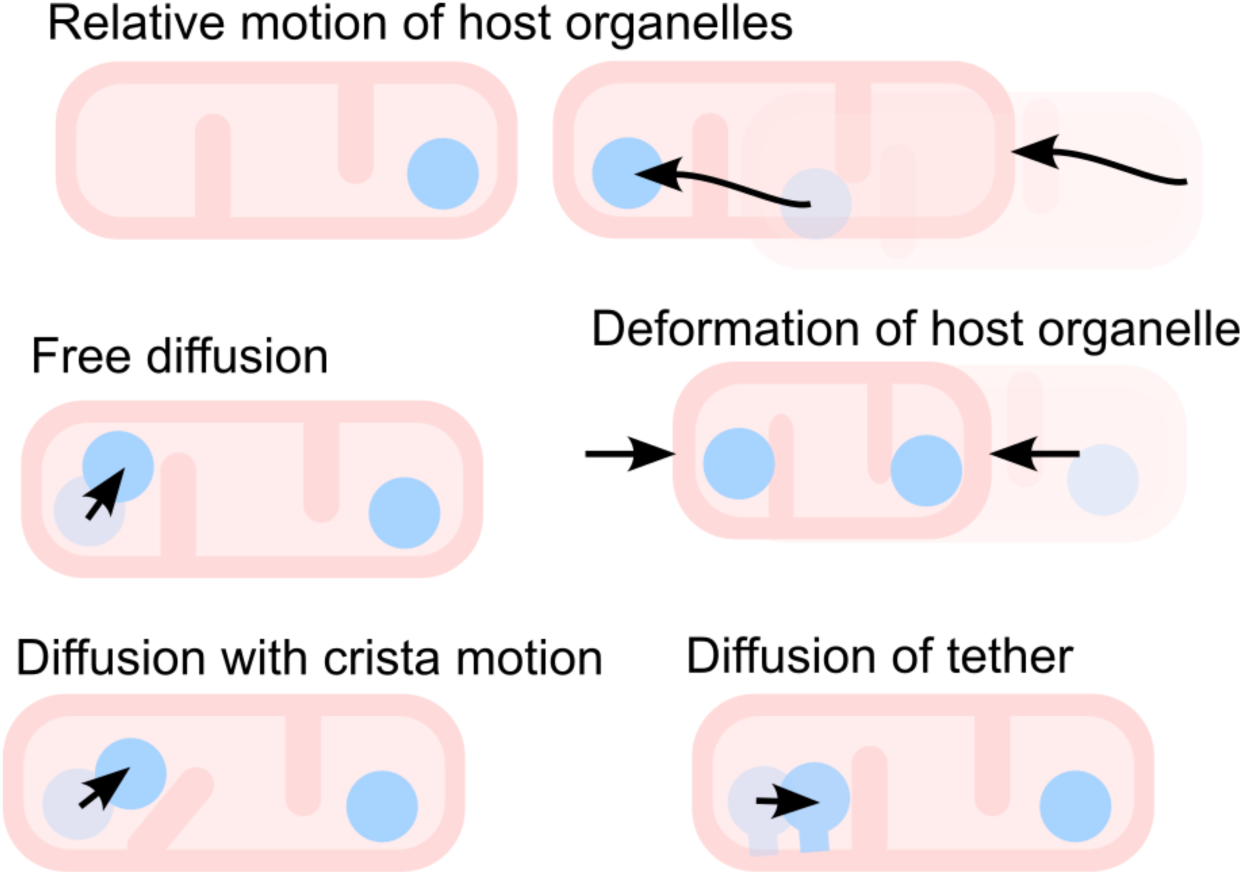
Different mechanisms by which relative motion of nucleoid pairs can be observed.

**Supplementary Figure 2.**
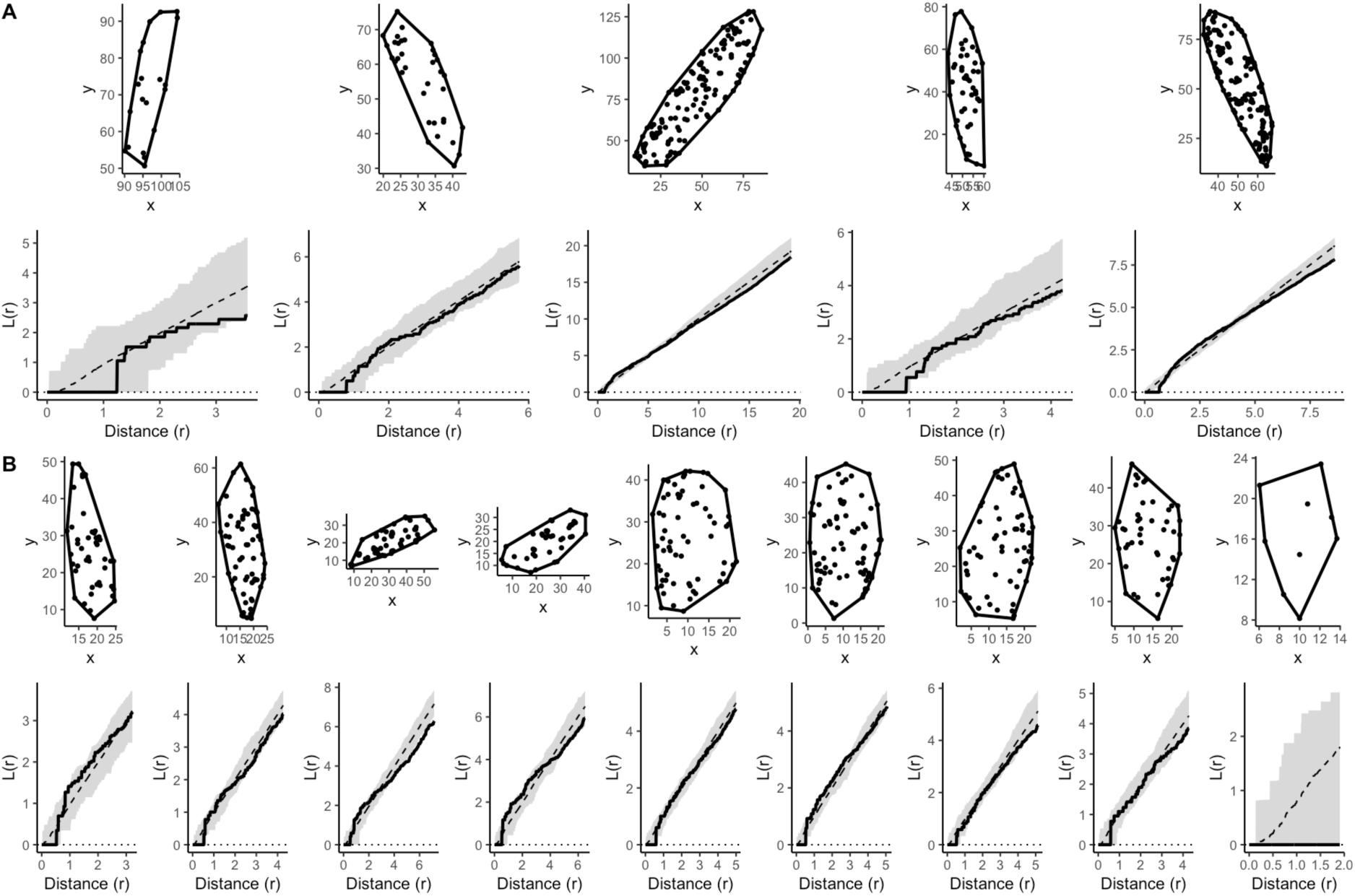
Spatial statistics for mitochondria (A) and nucleoids (B). Top rows show example distributions and convex hulls used in statistical modelling; bottom rows show comparison of empirical neighbour density (black) with distribution under the null model (grey).

**Supplementary Figure 3.**
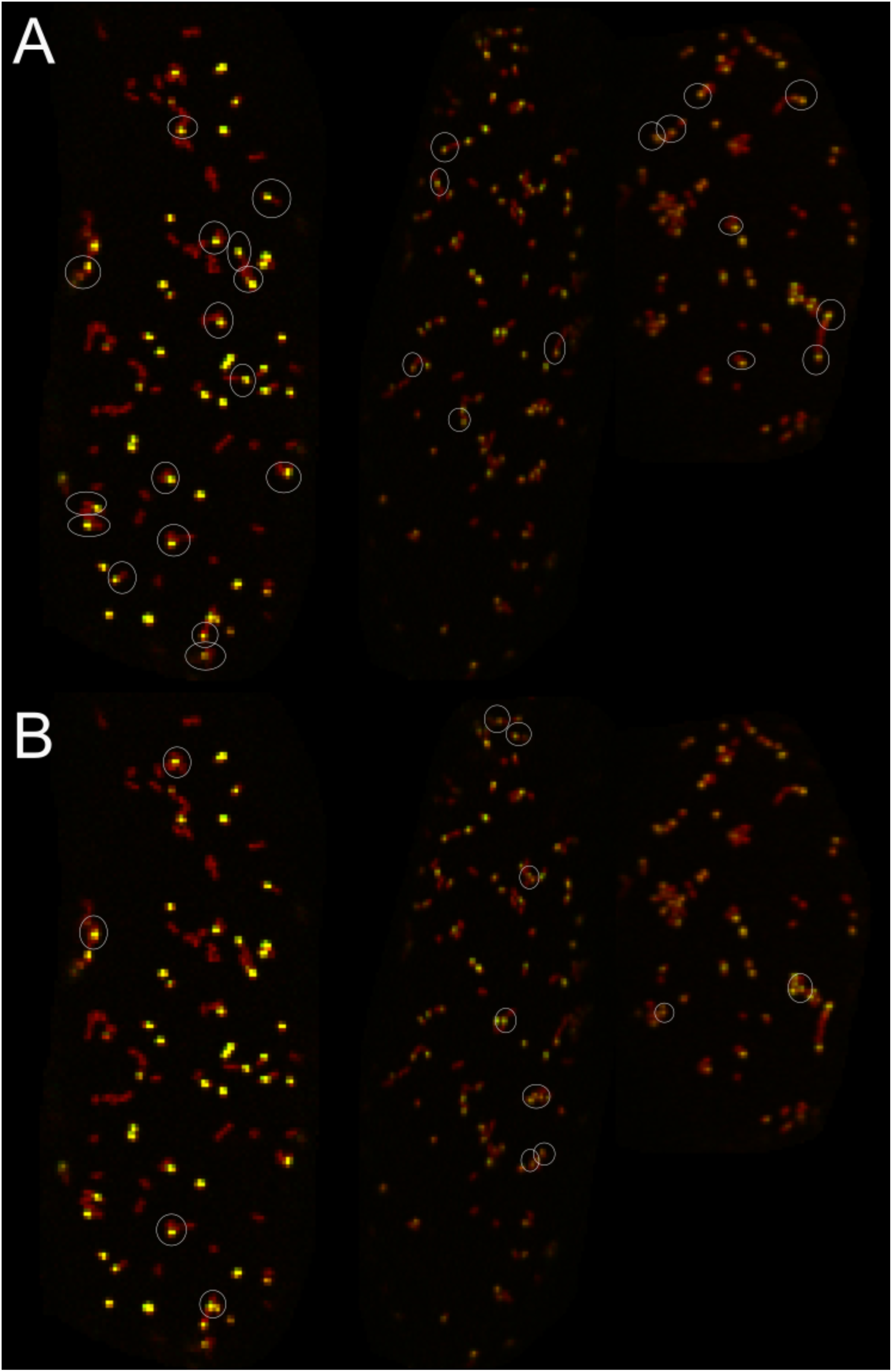
Enrichment of nucleoids at termini of elongated mitochondria. (A) Instances of nucleoid signal appearing at the end of an elongated mitochondria (≥ 6 px). (B) Instances of nucleoid signal appearing in the centre of an elongated mitochondria, with observable matrix signal either side of the nucleoid.

**Supplementary Figure 4.**
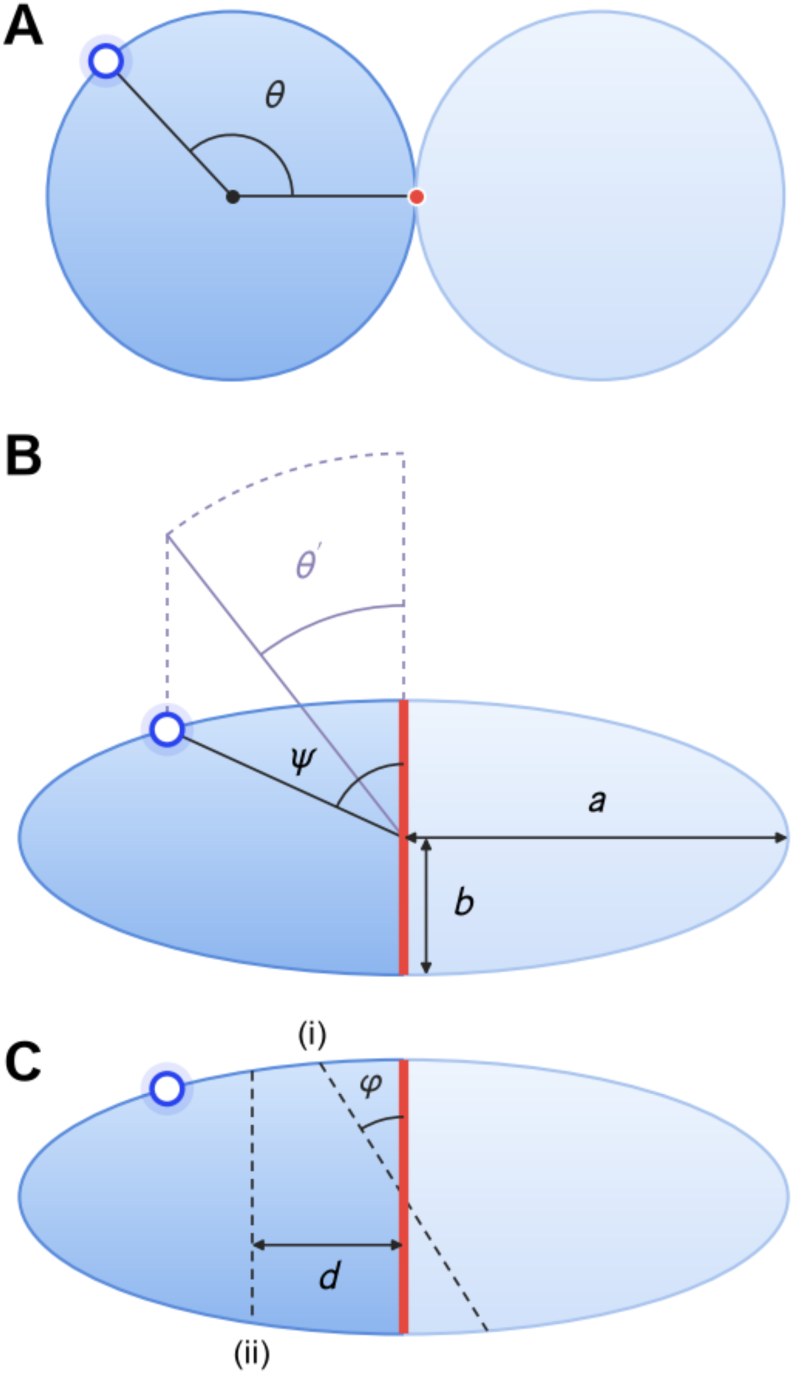
Geometry of fusion-fission model. (A) Two circular mitochondria about to fuse at the red point, with a nucleoid at angular position *θ*. (B) Elliptical fused mitochondrion, where the fusion point has expanded to the interface line (red) and total perimeter has been conserved. The nucleoid is located by its parametric angle *θ′* (grey construction), and by the geometric angle *ψ* between the fusion plane and the line joining the centre to the nucleoid, with tan 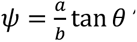. C) Two options for fission. (i) Fission plane runs through the centre of the ellipse, with angular difference *φ* from the fusion plane. (ii) Fission plane runs parallel to fusion plane, with horizontal offset *d*.

## Notes

https://github.com/StochasticBiology/mtdna-motion

### Competing Interest Statement

The authors have declared no competing interest.

